# Draft genome of the marine *Entamoeba* species reveals reduction in the gene family repertoire associated with pathogenicity and lateral gene transfer for adaptation to the marine environment

**DOI:** 10.1101/2025.02.12.637563

**Authors:** Tetsuro Kawano-Sugaya, Shinji Izumiyama, Tomoyoshi Nozaki

## Abstract

*Entamoeba* is the amoebozoan parasite commonly found in the intestines of animals. *E. marina* is the first exception isolated from marine sediments, possibly adapting from animal intestines to the sea. However, the evolutionary process of *E. marina* remains uncertain due to the lack of a genome sequence. Here, we present the *de novo* genome and transcriptome of *E. marina* using Oxford Nanopore MinION and Illumina HiSeq/MiSeq. The genome of *E. marina* is approximately 37.5 Mbp in length and consisted of 202 contigs, which is the second longest followed by *E. invadens*. *E. marina* showed significant reduction in the major virulence-associated gene families, including cysteine proteases, lysosomal enzyme transporters, and surface galactose/N-acetylglucosamine-specific lectins, suggesting diversification, more specifically reduction of pathogenicity-related genes. Genome and RNA-seq analyses also indicated genes either conserved throughout eukaryotes or laterally transferred from prokaryotes, and potentially responsible for salt tolerance. Our study provides insights into the mechanism underlying the lifestyle changes in the evolution of parasitic eukaryotes.

## Introduction

Acclimation to environmental changes is one the major evolutionary driving factors (Lamichhaney et al. 2015). *Entamoeba* is the genus that belong to the super group Amoebozoa and composed of mostly parasitic organisms that reside in a broad range of environments including the intestine of vertebrates of fish (*E. chiangraiensis*; Jinatham et al. 2019), reptilian (*E. invadens*; Meerovitch 1958), and non-vertebrates such as cockroaches (Kawano et al. 2017), as well as the oral cavity of humans (*E. gingivalis*; Smith and Barrett 1915). There are also a few free-living exceptions of *Entamoeba* that inhabit sewage water (*E. moshkovskii* Tshalaia 1941; see Scaglia et al. 1983) and marine sediments (*E. marina*; Shiratori and Ishida 2016). Previous phylogenetic studies indicated that *E. marina* diverged together with the intestinal *Entamoeba* species at the base of the *Entamoeba* clade (Shiratori and Ishida 2016; Kawano et al. 2017). This phylogenetic evidence is consistent with the premise that *E. marina* originally inhabited the intestines of animals, and then has subsequently adapted to the marine environment. However, it remains elusive whether *E. marina* is parasitic with the host remaining unidentified or it represents another free-living *Entamoeba* sibling.

The genomes of various parasitic and free-living *Entamoeba* have been sequenced (Loftus et al. 2005; Ehrenkaufer et al. 2013; Tanaka et al. 2019; Kawano-Sugaya et al. 2020; Nakada-Tsukui et al. 2018). The genome information of *Entamoeba* species that live in the conditions clearly divergent from other *Entamoeba,* such as *E. marina,* should indicate key genes or pathways necessary for the adaptation to such environments, i.e., sea water or sediments, or maybe a free-living lifestyle. Here, we describe the first draft genome assembly of the *E. marina* strain SRT209 (Shiratori and Ishida 2016), which consists of approximately 37.5 Mbp and encodes 10,771 proteins with 46% being hypothetical proteins. We found a reduction in the repertoire of virulence-associated gene families in *E. histolytica,* as well as diversification of genes related to vesicular transport and oxidative stress management. Furthermore, we identified a pair of genes, one conserved throughout eukaryotes and the other laterally transferred, that are potentially responsible for salt tolerance in *E. marina*. Taken together, a newly disclosed *E. marina* genome supports the premise that a combination of such genetic alterations plays a role for adaptation to diverse living environment and/or for transition from the parasitic to free-living lifestyle.

## Materials and Methods

### Organism and culture

*Entamoeba marina* strain SRT209 was kindly provided by Dr. Seiki Kobayashi (National Institute of Infectious Diseases) and cultivated using protocols modified from a previous publication (Shiratori and Ishida 2016). Briefly, the cells were cultured under monoxenic conditions with *Shewanella* sp. in 10% YIMDHA-S broth (Suzuki et al. 2008) diluted with artificial seawater (Cat. 395-01343; Nihon Pharmaceutical, Japan) at 25.5°C and passaged weekly.

### Short read genome sequencing

A total of 10^8^ trophozoites grown in 15 mL polystyrene non-treated culture tubes (CLS430157; Corning, AZ, USA) in the late logarithmic phase were harvested by centrifugation at 400 × g for 5 min. Total genomic DNA was isolated and purified using the QIAGEN Blood & Cell Culture DNA Kit with Genomic-tip 100/G (Qiagen, Germany). The purity and quantity of DNA samples were estimated using a NanoDrop 1000 (Thermo Fisher Scientific, MA, USA) and Qubit dsDNA HS assay kit (Thermo Fisher Scientific), respectively. Purified DNA was used to construct 8 kb and 20 kb long jump distance mate- pair genomic libraries and sequenced using the HiSeq 2000 platform (100MP; Illumina, CA, USA) by Eurofins Genomics (Tokyo, Japan). Finally, the 8 kb mate-pair sequencing produced a total of 115.2 million reads (10.2 Gbp), and the 20 kb mate-pair sequencing produced a total of 61.1 million reads (5.5 Gbp).

### Long read genome sequencing

Trophozoites were grown in 50 tubes (15 mL polystyrene non-treated culture tubes) and harvested by centrifugation at 400 × g for 5 min. High molecular weight DNA was isolated and purified using the NucleoBond HMW DNA Kit (Macherey-Nagel, Düren, Germany). The purity and quantity of the DNA samples were assessed using a NanoDrop 1000 (Thermo Fisher Scientific) and a Qubit dsDNA HS assay kit (Thermo Fisher Scientific), respectively. The purified DNA was used to prepare the library using a Rapid Sequencing Kit (SQK- RAD004; Oxford Nanopore Technologies, Oxford, UK). Sequencing was performed on the MinION platform using R9.4.1 Flow Cells (Oxford Nanopore Technologies) using MinKNOW 3.6.5. Base calling was performed using Guppy 3.2.10 (Oxford Nanopore Technologies). Finally, 435,782 reads (7.3 Gbp) were obtained.

### RNA-seq

Total RNA was isolated from log-phase trophozoites of *E. marina* SRT209 grown in 15 mL polystyrene non-treated culture tubes using TRIzol reagent. RNA (8.18 ng) was used to prepare the RNA-seq library using the ScriptSeq v2 RNA-seq Library Preparation Kit (Epicentre, WI, USA) following the manufacturer’s protocol. Sequencing was performed at the Pathogen Genomics Center (National Institute of Infectious Diseases, Tokyo, Japan) on the MiSeq platform (300PE; Illumina), yielding a total of 13.9 million reads (5.6 Gbp). The reads were trimmed using fastp v0.22.0 (Chen et al. 2018) with the following options (-q 30 - w 2). Read mapping was performed using HISAT2 with --max-intronlen 3000 (Kim et al. 2019). After removing reads corresponding to the rRNA region (see Supplementary Data) using intersectBed in BEDTools (Quinlan and Hall 2010), the transcripts per million (TPM) values were calculated using the TPMCalculator (Vera Alvarez et al. 2019).

### *De novo* assembly

The long reads from the MinION were trimmed using Nanofilt v2.8.0 (De Coster et al. 2018) with the following options (-q 8 --headcrop 50 -l 5000). The trimmed reads were then assembled using minimap2 v2.24 (Li 2018) and miniasm v0.3 (Li 2016). The assembly fasta file was generated using the awk command (awk ’/^S/{print ">"$2"\n"$3}’ | fold). The two contigs with too high (>50%) GC content compared to the *Entamoeba* genome out of 66 contigs were manually discarded as contamination from the co-cultured bacteria *Shewanella* sp. To correct misassemblies, the 64 contigs were divided into a total of 564 contigs using Tigmint v1.2.9 (Jackman et al. 2018) with subsequent options (tigmint-long span=auto G=37295306 dist=auto). The consensus sequence was called by Racon v1.5.0 (Vaser et al. 2017) for five times. The polishing step was performed using Pilon v1.24 (Walker et al. 2014) with the following options (-Xmx40G --tracks --verbose) using the short reads generated from Illumina mate-pair sequencings and trimmed by fastp. Final assembly had a total length of 37,460,168 bp consisted of 202 contigs. Assembly statistics were calculated using QUAST v5.0.0 (Mikheenko et al. 2018). The scripts used in this study are provided in the supplementary data.

### Gene prediction and annotation

We detected repeat regions in the assembled *E. marina* genome using RepeatModeler v2.0.1 (Flynn et al. 2020) and soft-masked them using RepeatMasker v4.1.1 (Smit et al. 2013; http://www.repeatmasker.org). Gene prediction was performed by BRAKER2 (Brůna et al. 2021), providing OrthoDB v10 (Kriventseva et al. 2019) as a protein database. Predictions with supports based on OrthoDB were used for the subsequent process. The remainder of the prediction was additionally filtered with RNA-seq data. The RNA-seq reads from MiSeq were assembled by Trinity v2.15.1 (Grabherr et al. 2011). The reads were remapped to the RNA-seq assembly by HISAT2 (Kim et al. 2019). The reads mapped on RNA-seq were recovered from bam file and then mapped to the genome assembly using HISAT2 with -- max-intronlen option. The annotation of related species *Entamoeba histolytica* HM-1:IMSS provided in AmoebaDB release 63 (2022-12-06; Aurrecoechea et al. 2011) was transferred to *E. marina* genome using pipelines in GeMoMa 1.9 (Keilwagen et al. 2019). Annotations from GeMoMa and BRAKER2 were merged by GffCompare v0.12.6 (Pertea and Pertea 2020) using the annotation from GeMoMa as a reference. As a result, a total of 10,771 genes were identified. They were compared to the UniProt Reference Clusters (UniRef90; Suzek et al. 2015) in EnTAP v0.10.8 (Hart et al. 2020). Finally, total a total of 9,363 genes (including 4,954 hypothetical or uncharacterized genes) were annotated (Supplementary Data Table S1). Statistics for genes were calculated by AGAT (https://www.doi.org/10.5281/zenodo.3552717).

### Phylogenetic analysis

Sequences in *Entamoeba* species were retrieved from AmoebaDB release 63 using BLASTp (threshold: 1e^-5^). They were aligned using the mafft-linsi command in MAFFT v7.520 (Katoh and Standley 2013). TrimAl v1.2 (Capella-Gutiérrez et al. 2009) automatically selected well-aligned positions using the -automated 1 option. Phylogenetic analyses were performed using FastTree 2.1.10 with the -lg option (Price et al. 2010). The results were visualized using FigTree v1.4.4 (https://github.com/rambaut/figtree).

### Identification of Rab GTPases in *E. marina*

When we used BLASTp to identify Rab GTPases in the *E. marina* genome using the criteria from a previous study (Saito-Nakano et al. 2005), it gave innegligible false positives from other Ras superfamilies due to GTPase domains. Thus, we conducted RPS-BLAST (Altschul et al. 1997) searches using proteome in *E. marina* as queries and conserved domains database (CDD) as a database to evaluate if Rab GTPases from *E. marina* under analysis most likely contain Rab GTPase conserved domains (PSSM-ID: 206640, 206653, 206654, 206655, 206656, 133267, 206657, 206658, 206659, 206660, 206661, 206688, 206689, 133306, 206692, 206693, 206694, 133310, 133311, 206695, 206696, 133314, 133315, 206697, 206698, 133318, 133319, 206699, 133321, 133322, 133323, 133324, 133326, 206700, and 206701; Table S4). We defined Rab GTPases based on top-hits under the e-value 1e^-10^.

## Results and Discussion

### *E. marina* possesses the second largest genome in *Entamoeba,* but its gene repertoire resembles other parasitic *Entamoeba*

The draft genome of *E. marina* is the second largest among available *Entamoeba* genomes, with a total length of 37,460,168 bp, and consisted of 202 contigs. This places it second in size, following *E. invadens* (40.9 Mbp), among the sequenced *Entamoeba* species including *E. histolytica*, *E. dispar*, *E. moshkovskii*, and *E. invadens* (Fig. 1A). The basic genomic characteristics of *E. marina*, including GC content and gene number, fall within the range observed in other *Entamoeba* species (Table 1). The *E. marina* genome encodes 10,771 proteins, of which 4,954 are annotated as hypothetical proteins.

**Figure 1.**
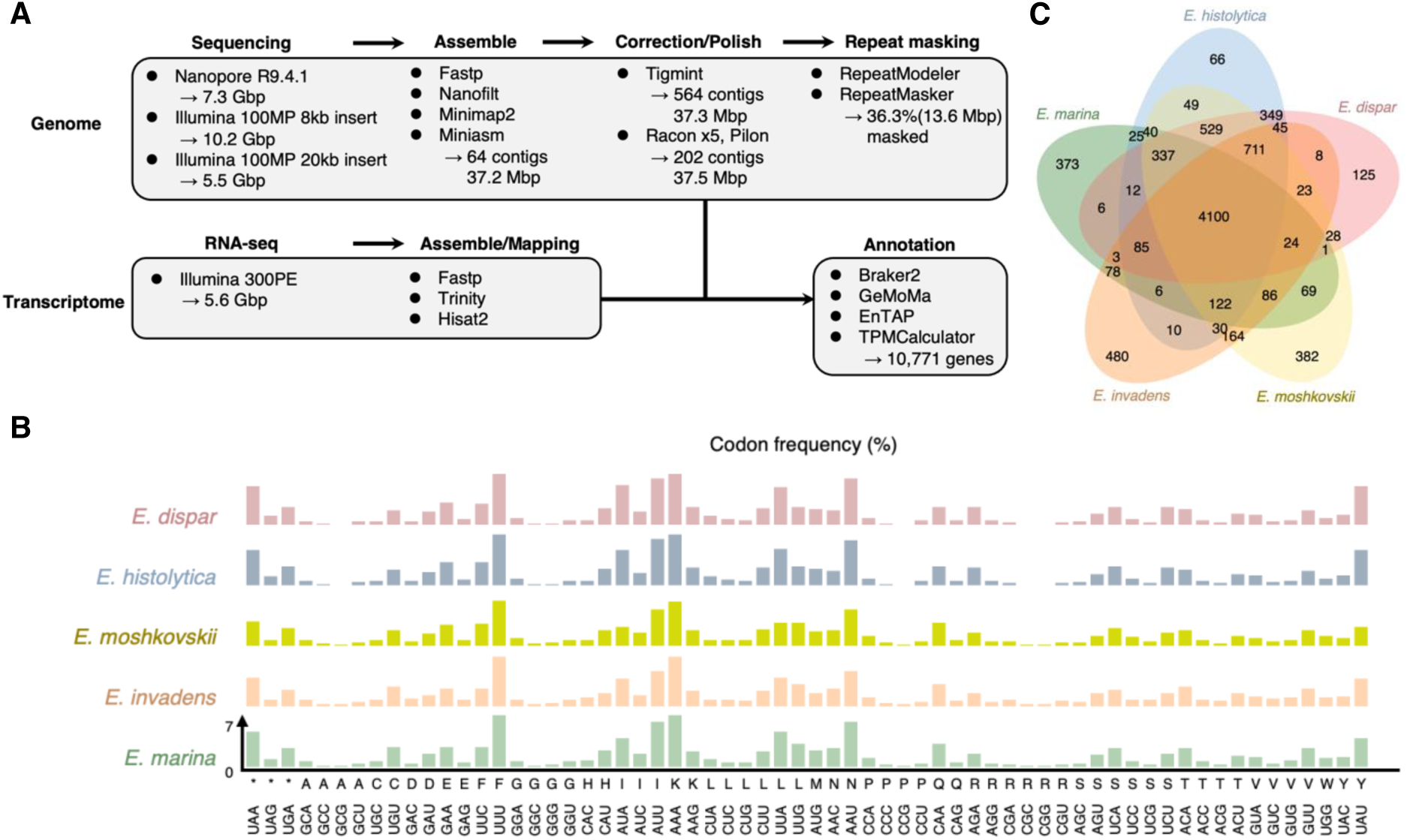
Similarities among the genomes of *E. marina* and other *Entamoeba*. (*A*) The flow diagram for genome analysis of *E. marina*. (*B*) Codon usage in *Entamoeba* genomes. The y-axis represents the frequency (%) of each codon. (*C*) Venn diagram showing the inclusion relation of the orthogroups in *Entamoeba*.

**Table 1.** Basic features of the genome assemblies of five *Entamoeba* species. Statistics for genome assemblies of *Entamoeba*. The data for *E. dispar*, *E. histolytica* HM-1:IMSS, *E. moshkovskii*, and *E. invadens* were retrieved from AmoebaDB.org. The data for *E. histolytica* HM-1:IMSS Clone 6 2001 was from our previous study (Kawano-Sugaya et al. 2020).

A notable feature of the *E. marina* genome is its high repeat contents, with repeat elements accounting for 36.3% of the genome, as detected by RepeatModeler and RepeatMasker. This repeat-rich nature of the *E. marina* genome is similar to *E. histolytica* HM-1:IMSS Clone 6 2001 (37.6%; Kawano-Sugaya et al. 2020). Furthermore, the codon usage and overall repertoire of *E. marina* are comparable to those of other *Entamoeba* species (Fig. 1B; Table S1). Orthologous clusters identified by OrthoVenn2 (Xu et al. 2019) revealed 4,100 conserved clusters across five *Entamoeba* species, with 373 clusters unique to *E. marina* (Fig. 1C; Table S3). These findings suggest that, despite its large genome, *E. marina* maintains a gene repertoire similar to other parasitic *Entamoeba* species, indicating that its evolutionary divergence may not be strongly associated with the expansion of virulence-related gene families.

### Reduction in the repertoire of virulence-associated gene families in *E. marina*

We investigated key virulence-associated genes in *E. marina*, including cysteine proteases (CP), cysteine protease-binding protein family (CPBFs; Marumo et al. 2014), intrinsic inhibitor of CPs (ICP; Sato et al. 2006), and Rab small GTPases, using BLAST (Altschul et al. 1990). CPs play critical roles in the pathogenesis of *E. histolytica*, such as cell killing, phagocytosis, trogocytosis, destruction of extracellular matrix and tissues (Irmer et al. 2009; Ralston et al. 2014), and encystation (Ebert et al. 2008), which is necessary for human-to-human transmission. Our BLAST search revealed that approximately half of the CP family A members present in other human-infecting *Entamoeba* species are absent in *E. marina* (Fig. 2A; CP-A1, A2, A4, A7, A9, A11, and A12; labeled in white). On the other hand, *E. marina* possesses a species-specific subfamily of CP family A genes (cyan; Fig. 2A), suggesting that these unique CPs may have specialized biological roles in *E. marina*.

**Figure 2.**
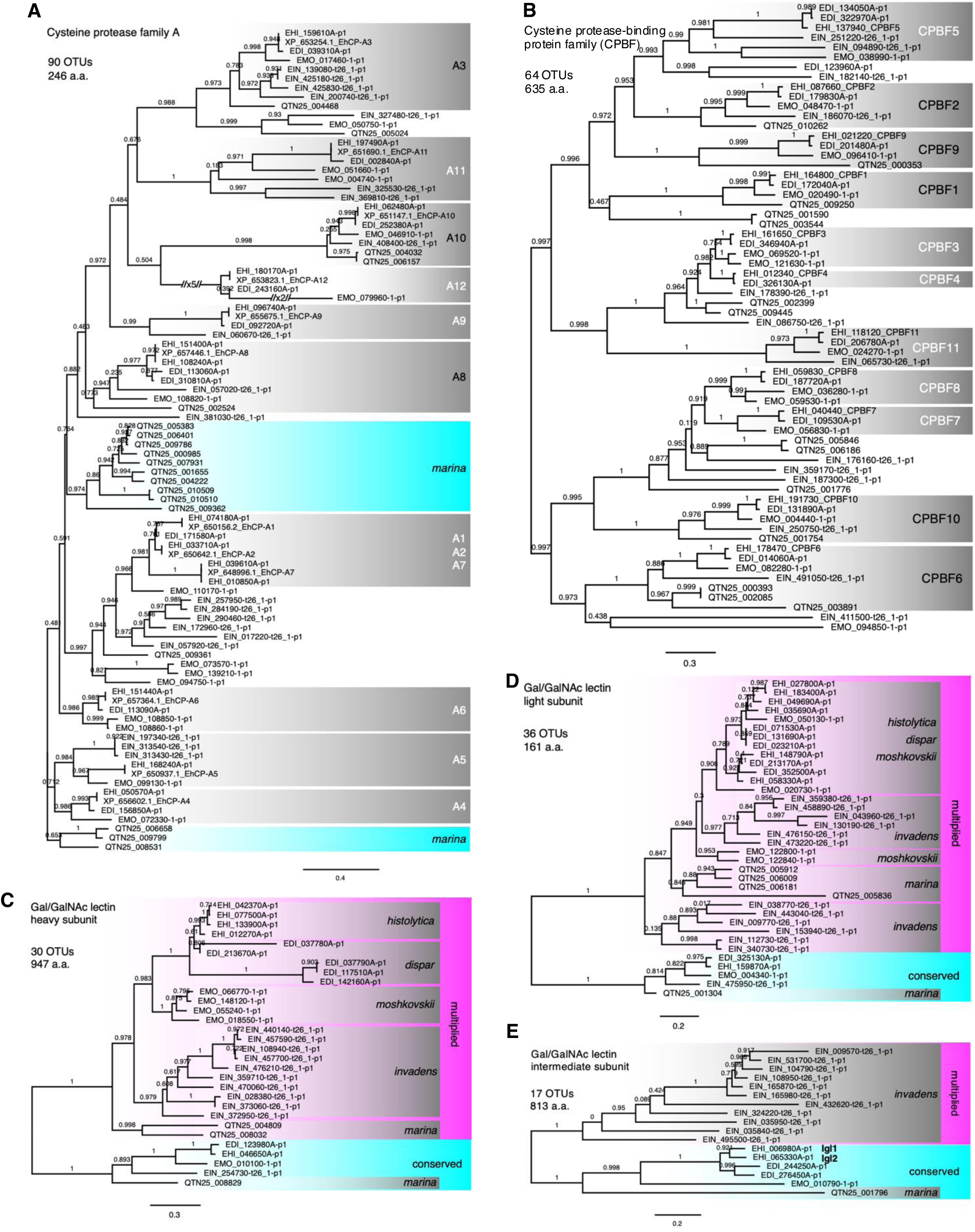
Half-loss of virulence-associated genes in *E. marina*. The maximum likelihood tree of virulence-associated gene families in the five *Entamoeba* including (*A*) cysteine protease family A genes, (*B*) cysteine protease-binding protein families, (*C*) lectin heavy subunits, (*D*) lectin light subunits, and (*E*) lectin intermediate subunits. The bootstrap values are shown on each branch. The absence or presence of subgroups in *E. marina* are shown in white or black, respectively.

Intracellular traffic and secretion of lysosomal enzymes, including CPs, are regulated by specific transporters such as CPBF1 (Marumo et al. 2014; Nakada-Tsukui et al. 2020) and inhibitors like ICPs (inhibitors of cysteine proteases). The repertoire of CPBPs in *E. marina* was markedly different from that of other *Entamoeba* species. Specifically, six of the eleven CPBFs (CPBF3, 4, 5, 7, 8, and 11) were absent in *E. marina*, while the other CPBFs (CPBF1, 2, 6, 9, and 10) were conserved (Fig. 2B; labelled in white). Among them, CPBF1 is the only CP carrier (receptor) among the CPBF members, responsible for transporting all family A CPs from the endoplasmic reticulum to lysosomes (Marumo et al. 2014). ICPs are intrinsic proteins that bind CPs, inhibit CP activities, and prevent their secretion (Sato et al. 2006). In terms of ICPs, *E. marina* retained two ICPs (QTN25_002404 and QTN25_005631), based on a BLASTp search against ICP1 of *E. histolytica* (EAL47869.1), showing 35.9% and 25.7% identities, respectively. These values are comparable to those observed between *E. histolytica* ICP1 and its *E. invadens* orthologs (XP_004185788.1 and XP_004256329.1) (48.5% and 27.5%). Notably, CPBF6 and CPBF8 regulate the transport of lysosomal hydrolases involved in glycoprotein and carbohydrate degradation in *E. histolytica* (Furukawa et al. 2012; Furukawa et al. 2013). In *E. histolytica,* it was demonstrated that CPBF6 binds to α-amylase and γ-amylase while CPBF8 binds to lysozymes and β-hexaminidase. In *E. marina,* only CPBF6, is retained, while CPBF8 is absent. This absence is particularly interesting because CPBF8 is also missing in *E. invadens*, suggesting that CPBF8 was likely acquired during the divergence of the common ancestor of *E. histolytica, E. dispar,* and *E. moshkovskii* from the lineage leading to *E. marina* and *E. invadens*.

A similar pattern of the absence of isotypes in *E. marina* may apply to CPBF3 and CPBF4, as the clade containing *E. marina* and *E. invadens* is positioned at the base in the phylogenetic tree relative to the CPBF3/4 clade. In contrast, CPBF5 and CPBF11 are absent in *E. marina,* but are retained in *E. invadens*, suggesting lineage-specific expansions of certain CPBF isotypes after the divergence of *E. marina* from other *Entamoeba* species. Note that it was demonstrated that CPBF3 and CPBF4 likely makes heterodimeric complex and the ligands of CPBF3, 4, 5, and 11 remain undermined.

Adherence to and cytolysis of the intestinal epithelia are hallmarks of *E. histolytica* pathogenesis. While adherence, but not cytolysis, is conserved across *Entamoeba* species, including commensal, non-invasive siblings, *E. histolytica* uses the surface galactose/N-acetylgalactosamine inhibitable lectin as a receptor for adhesion to the host epithelium. The Gal/GalNAc lelcting of *E. histolytica* consists of three subunits, heavy (Hgl), intermediate (Igl), and light subunits (Lgl), whose encoding genes are conserved across the *Entamoeba* genus (Frederick and Petri 2005). Hgl is a transmembrane protein with the amino-terminal carbohydrate-recognition domain (LecA), responsible for binding to specific carbohydrate epitopes on the human target cells, while Lgl is GPI-anchored and forms a disulfide bond with Hgl. Igl, also GPI anchored, plays a role in adherence (Petri et al. 2006; Min et al. 2016; Kato and Tachibana 2022). Phylogenetic analysis of Hgl and Lgl across *E. marina* and four other *Entamoeba* revealed similar topologies (Fig. 2C-D). Both Hgl and Lgl trees consist of two major clades: the first clade contains the core cognate members from each *Entamoeba* species (cyan), while the second clade includes expanded members (magenta). This pattern suggests an early gene duplication event in the common ancestor, followed by the additional gene duplication leading to species-specific subclades (magenta). In contrast, Igl showed no diversification across *Entamoeba* species, with the exception of *E. invadens*, which has 11 isoforms (Fig. 3E; magenta).

**Figure 3.**
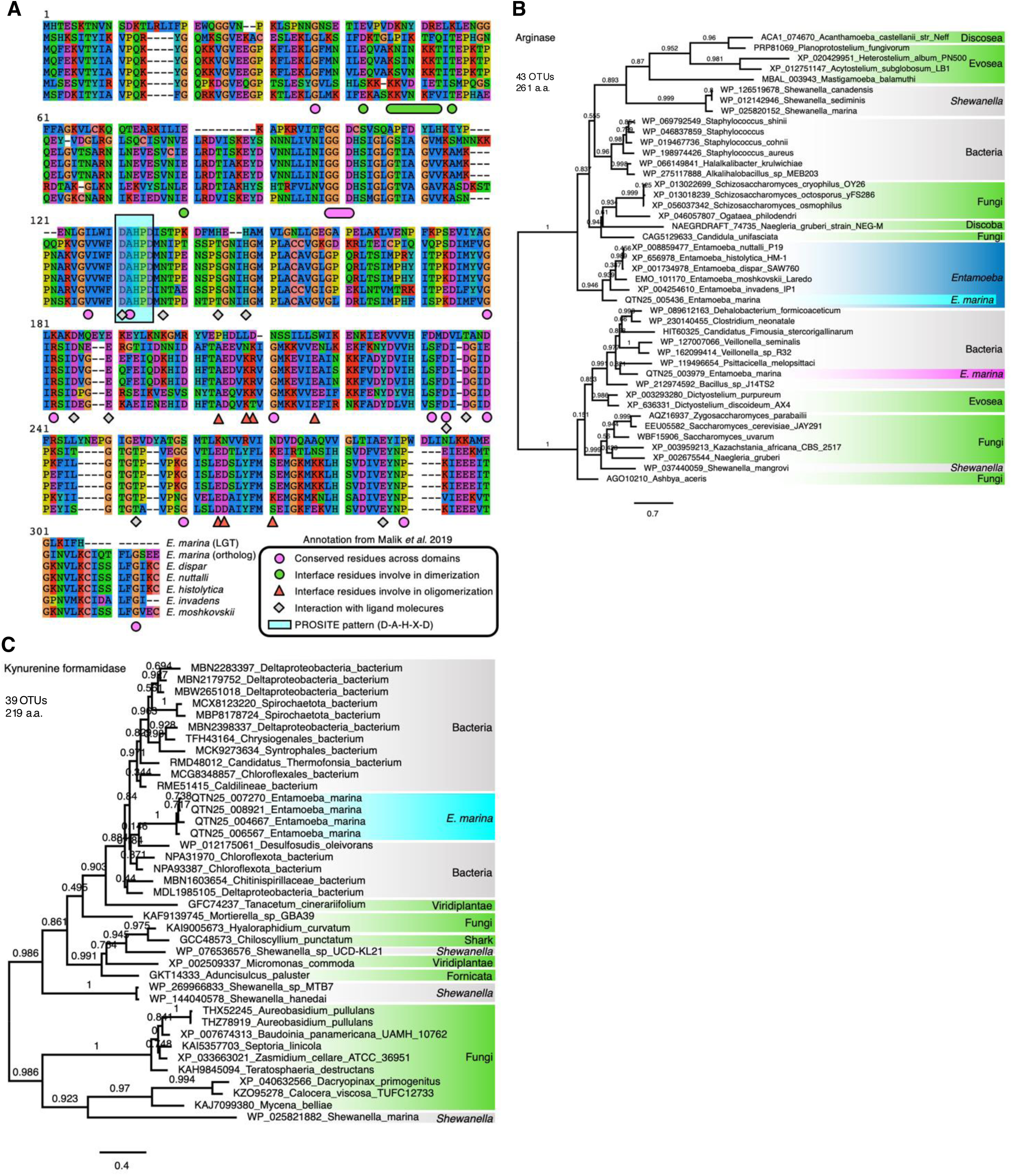
The genes potentially related to the adaptation to the marine environment. (*A*) The sequence alignment of arginases from *E. marina* and other *Entamoeba*. (*B*) The maximum likelihood tree of arginases from *E. marina*. The orthologous arginase in *Entamoeba* is shown in cyan, whereas the *E. marina*-specific arginase is shown in magenta.

### Lineage-specific diversification of major gene families in *E. marina*

The diversification and expansion of gene families through duplication has been well documented in various gene families including those encoding *Bacteroides* surface protein A (BspA) family (Lorenzi et al. 2010), avrRpt2-induced gene 1 (AIG1) family (Nakada-Tsukui et al. 2018), and Rab GTPases (Saito-Nakano et al. 2005). These gene families are known to be associated with the pathogenicity of *Entamoeba*. Our analysis revealed that *E. marina* lack the AIG1 family, a feature shared with *E. invadens* and *E. moshkovskii*. In contrast, the BspA family is highly diversified in *E. marina*, which encodes a total of 449 BspA proteins, approximately four times more than the 121 BspA proteins found in *E. histolytica*. (Table S1).

Rab small GTPases, conserved across eukaryotes, regulate directional vesicular traffic between cellular compartments in a GTP- or GDP-dependent manner. The genomes of *Entamoeba* species contain a large number of Rab GTPases, indicating extensive diversification of this protein family in the genus (Table S5). The number of Rab GTPases (111-142 genes per species) exceeds that in multicellular eukaryotes, including humans (∼70), suggesting that Rab GTPases play a complex role in regulating membrane traffic in *Entamoeba* (Saito-Nakano et al. 2005). Notably, many Rabs (e.g., RabA, RabB, Rab5, Rab7A, Rab7B, Rab7D, Rab8A, Rab11A, Rab11B, Rab11D, Rab21, Rab35) are involved in key virulence and transmission-related processes such as phagocytosis, trogocytosis, and encystation (Saito-Nakano et al. 2004; Welter et al. 2005; Mitra et al. 2007; Hernandes-Alejandro et al. 2013; Emmanuel et al. 2015; Verma and Datta 2017; Saito-Nakano et al. 2021). Among the five *Entamoeba* species analyzed, *E. marina* has the highest number of Rab small GTPases, encoding a total of 160 proteins for 76 subfamilies (e.g., Rab7) (Table S5), based on RPS-BLAST search. Many gene duplications were identified specifically in *E. marina*, contributing to the expansion of the Rab GTPase family. Of 76 subfamilies, twenty include multiplicated isotypes, including Rab7D, Rab7H, Rab7I, Rab11C, RabA, RabC1, RabC7, RabD1, RabF5, RabK3, RabM1, RabM3, RabP2, RabX6, RabX11, RabX14, RabX15, RabX23b, RabX33 and RabX33b.

TLDc [Tre2/Bub2/Cdc16 (TBC), Lysin motif (LysM), Domain catalytic] domain- containing proteins are another gene family that shows extensive diversification in *Entamoeba*. The *E. marina* genome encodes a total of 246 TLDc domain-containing proteins (Table S1), which is 2.1- to 5.7-fold more than those in other parasitic *Entamoeba* species (43-62 in *E. histolytica*, 50 in *E. dispar*, 50 in *E. nuttalli*, and 116 in *E. invadens*), but less than the number in the free-living *E. moshkovskii* (499). Proteins containing a TLDc domain are known to provide oxidative resistance by reducing reactive oxygen species (Finelli et al. 2016) and to interact with proton-pumping V-ATPase (Eaton et al. 2021).

### Genes potentially related to lifestyle conversion in *E. marina*

In an effort to identify genes responsible for lifestyle conversion to the marine environment, we hypothesized that the genes gained by *E. marina* via lateral gene transfer (LGT) might have played significant role in this process. We investigated potential LGT genes that are exclusively present in *E. marina* but absent or nearly absent in other *Entamoeba* species.

According to annotations from EnTAP (Hart et al. 2020), we identified a total of 95 bacterial-origin genes gained by *E. marina* via LGT. Of these, eighty two were also shared by at least one other *Entamoeba* species, while thirteen genes were unique to *E. marina,* making them strong candidates for LGT-derived genes involved in marine adaption (Table 2).

**Table 2.** *E. marina* specific LGT genes. Sequence identifier, length, TPM, hit in UniRef90, EggNOGTaxScope, and EggNOGDescription for the 13 genes that *E. marina* specifically acquired via lateral gene transfer.

Our RNA-seq analysis confirmed that among 13 LGT genes, several – such as genes encoding one arginase (QTN25_003979) and four cyclase-like (QTN25_004667, QTN25_006567, QTN25_007270, and QTN25_008921) were expressed at levels exceeding the median value of all transcripts (TPM > 7.101; Table S6). In addition to the LGT-acquired arginase, *E. marina* also contains a gene encoding a conserved arginase that is found in all *Entamoeba* species. Interestingly, neither the *E. marina-*specific LGT-derived arginase nor the conserved *Entamoeba* arginase contain an organelle-targeting transit peptide, unlike mitochondria-targeted arginase-2 in humans. Alignment of amino acid sequences revealed a highly conserved PROSITE pattern (D-A-H-X-D) across all *Entamoeba* arginases (Fig. 3A; cyan), and residues involved in ligand interactions are also well conserved (Fig. 3A; grey diamond), consistent with previous studies (Malik et al. 2019). In contrast, residues involved in oligomerization (Malik et al. 2019) were not conserved in the *E. marina-*specific arginase, suggesting a potential lack of oligomerization or differences in the oligomerization state (Fig. 3A; triangle). Phylogenetic analysis indicated that the evolutionary origins of the *E. marina*- specific LGT-derived arginase and the conserved *Entamoeba* arginase were clearly distinct (Fig. 3B; magenta and blue rectangle). The close phylogenetic relationship between of *E. marina-*specific arginase and arginase from *Psittacicella melopsittaci*, a Gram-negative bacterium in γ−proteobacteria, indicates this *E. marina-*specific arginase gene was acquired from this bacterial lineage (Fig. 3B; magenta). In contrast, the other arginases common to *Entamoeba* did not show close kinship with other organisms (Fig. 3B; blue rectangle). These findings suggest that arginases may have undergone frequent replacement events throughout evolution, providing selectable advantages under unique habitat conditions (Fig. 3B; green).

Furthermore, the *E. marina* genome encodes four closely related proteins that show similarity to bacterial kynurenine formamidase (UniRef90_A0A7C3CCR2), with e-values ranging from 5.71e^-88^ to 6.48e^-107^ (52.8–61.3% similarities and 93.7–94.7% coverages; Table S1). These *E. marina* proteins also exhibited 53.97–60.08% similarit to cyclase from a *Deltaproteobacteria bacterium* (MBN2538769.1), with 94–95% coverage and e-values ranging from 4e^-91^ to 5e^-109^ as determined by in NCBI BLAST. Kynurenine formamidase catalyzes the hydrolysis of N-formyl-L-kynurenine to L-kynurenine, an intermediate in the kynurenine pathway of tryptophan degradation. L-kynurenine is further transaminated to kynurenic acid by kynurenine transaminase, which is known for its neuroprotective effects and antioxidant activity, scavenging free radicals originating from FeSO_4_ (Lugo-Huitrón et al. 2011). RNA-seq data indicate that these L-kynurenine formamidase genes (TPM = 198.5 ∼ 2,738.2; Table 2) and two kynurenine transaminase genes (TPM = 98.0 for QTN25_008178 and 70.3 for QTN25_008089; Table S6) are expressed in *E. marina*, suggesting their involvement in defense against oxidative stresses in trophozoites. The phylogenetic tree indicates that *E. marina* cyclase-like protein genes were apparently acquired from bacteria by LGT, with distinct bacterial origins compared to other eukaryotic cyclase-like genes (Fig. 3C). In addition to its antioxidant properties, kynurenine has been implicated in several other biological functions, including maintenance of NAD^+^ levels, as well as immune tolerance and inflammatory responses and neuroprotective effects. Since the kynurenine pathway ultimately contributes to the synthesis of NAD^+^, a critical molecule for cellular energy production and redox homeostasis, it is possible that *E. marina* might use this pathway not only to defend against oxidative stress but also to maintain cellular energy balance under the challenging marine conditions. In addition, well-documented neuroprotective effects of kynurenic acid, a metabolite of kynurenine, including its role in preventing neurodegenerative diseases may be relevant to the survival and functionality of *E. marina* under stress conditions in its marine habitat.

## Conclusion

Here, we presented a draft genome of ocean-derived *E. marina* and conducted genome comparisons between *E. marina* and other *Entamoeba* species, with aims to elucidate the mechanisms by which *E. marina* adapted to the marine environment. The *E. marina* genome largely resembles those of other *Entamoeba* species, with the majority of genes and gene families being conserved. However, we observed a reduction in the number or a loss of virulence-associated gene families. Notably, the loss of a large fraction of CPs and CPBFs is considered to be associated with lack of virulence to vertebrate and invertebrate hosts.

Distinct evolutionary patterns for three subunits for Gal/GalNAc lectins, known to play a pivotal role in recognition of and binding to the target organisms (pray), indicated that the lectins evolved in a way how individual species adapt to different environment. In addition, *E. marina* has undergone an expansion of certain gene families, including the BspA family, Rab family, and TLD genes, which are involved in the regulation of essential biological processes. On the contrary, repertoire expansion of AIG1 gene family, commonly observed in *Entamoeba* species that infect primates, such as *E. nuttalli* and *E. dispar*, did not occur in *E. marina*. This observation is consistent with the premise that AIG1 repertoire expansion contribute to virulence and mammalian host specificity. We also identified 13 genes uniquely acquired through LGT in *E. marina*, including genes encoding kynurerine formamidases and arginase. Taken together, the first draft genome of marine free-living *Entamoeba marina* demonstrated striking diversifications from other *Entamoeba* species, which underlie its unique features in marine adaptation and potentially free-living lifestyle or parasitism to organisms in marine sediments.

## Acknowledgements

We thank Seiki Kobayashi and Emi Sato-Ebine (National Institute of Infectious Diseases; NIID), Tetsuo Hashimoto (University of Tsukuba), and Kumiko Shibata (The University of Tokyo) for technical assistance on cultivation. We also thank Avik Kumar Mukherjee, Tsuyoshi Sekizuka, and Makoto Kuroda (NIID) for initial analysis of the *E. marina* genome. This study was supported in part by Grant for Research on Emerging and Re-emerging Infectious Diseases from Japan Agency for Medical Research and Development (AMED, JP22fk0108138 to TN), Grants-in-Aid for Scientific Research (B) (KAKENHI JP21H02723 to TN) from the Japan Society for the Promotion of Science (JSPS), and Grant for Science and Technology Research Partnership for Sustainable Development (SATREPS) from AMED and Japan International Cooperation Agency (JICA) (JP22jm0110022 to TN).

## Author contribution

TN conceptualized and supervised whole study and acquired funds. TKS determined methodology and investigated genome data. SI gave essential advice and technical supports. TKS and TN wrote the manuscript. All authors read and approved the final manuscript.

## Data availability

All sequence data produced in this study were deposited in the Sequence Read Archive (SRA) and GenBank at the National Center for Biotechnology Information (NCBI). The identifiers in the BioProject and BioSample are PRJNA985239 and SAMN35793876, respectively. The SRA accessions for the raw reads after quality trimming were as follows:1) SRR24958723 for long read genome sequence using Oxford Nanopore MinION; 2) SRR24958724 and SRR24958725 for genome sequence with 8kb or 20kb insert using Illumina 100MP sequencings; and 3) SRR24958722 for RNA-seq using Illumina 300PE sequencing. The final genome assembly and annotations were deposited in GenBank (accession number JAUKTS010000000). The all authors confirmed that all supporting data have been provided in the article through the supplementary data files.

## Supplemental Data

Supplemental files | Assembly and annotation data used in this study

Table S1. Annotation of *E. marina* genome from EnTAP

Table S2. Codon usage in *Entamoeba* with frequencies

Table S3. *E. marina*-specific orthologous clusters

Table S4. List of entries for Rab conserved domains in CDD

Table S5. Rab GTPases in *E. marina*, *E. histolytica*, *E. dispar*, *E. moshkovskii*, and *E. invadens*

Table S6. TPM for genes of *E. marina* in RNA-seq

## References

Altschul SF, Gish W, Miller W, Myers EW, Lipman DJ. 1990. Basic local alignment search tool. J. Mol. Biol. 215:403–410.

Altschul SF, Madden TL, Schäffer AA, Zhang J, Zhang Z, Miller W, Lipman DJ. 1997. Gapped BLAST and PSI-BLAST: a new generation of protein database search programs. Nucleic Acids Res. 25:3389–3402.

Aurrecoechea C, Barreto A, Brestelli J, Brunk BP, Caler EV, Fischer S, Gajria B, Gao X, Gingle A, Grant G, et al. 2011. AmoebaDB and MicrosporidiaDB: functional genomic resources for Amoebozoa and Microsporidia species. Nucleic Acids Res. 39:D612–9.

Brůna T, Hoff KJ, Lomsadze A, Stanke M, Borodovsky M. 2021. BRAKER2: automatic eukaryotic genome annotation with GeneMark-EP+ and AUGUSTUS supported by a protein database. NAR Genom Bioinform 3:lqaa108.

Capella-Gutiérrez S, Silla-Martínez JM, Gabaldón T. 2009. trimAl: a tool for automated alignment trimming in large-scale phylogenetic analyses. Bioinformatics 25:1972– 1973.

Chen S, Zhou Y, Chen Y, Gu J. 2018. fastp: an ultra-fast all-in-one FASTQ preprocessor. Bioinformatics 34:i884–i890.

De Coster W, D’Hert S, Schultz DT, Cruts M, Van Broeckhoven C. 2018. NanoPack: visualizing and processing long-read sequencing data. Bioinformatics 34:2666–2669.

Eaton AF, Brown D, Merkulova M. 2021. The evolutionary conserved TLDc domain defines a new class of (H+)V-ATPase interacting proteins. Sci. Rep. 11:22654.

Ebert F, Bachmann A, Nakada-Tsukui K, Hennings I, Drescher B, Nozaki T, Tannich E, Bruchhaus I. 2008. An Entamoeba cysteine peptidase specifically expressed during encystation. Parasitol. Int. 57:521–524.

Ehrenkaufer GM, Weedall GD, Williams D, Lorenzi HA, Caler E, Hall N, Singh U. 2013. The genome and transcriptome of the enteric parasite Entamoeba invadens, a model for encystation. Genome Biol. 14:R77.

Emmanuel M, Nakano YS, Nozaki T, Datta S. 2015. Small GTPase Rab21 mediates fibronectin induced actin reorganization in Entamoeba histolytica: implications in pathogen invasion. PLoS Pathog. 11:e1004666.

Finelli MJ, Sanchez-Pulido L, Liu KX, Davies KE, Oliver PL. 2016. The evolutionarily conserved Tre2/Bub2/Cdc16 (TBC), lysin motif (LysM), domain catalytic (TLDc) domain is neuroprotective against oxidative stress. J. Biol. Chem. 291:2751–2763.

Flynn JM, Hubley R, Goubert C, Rosen J, Clark AG, Feschotte C, Smit AF. 2020. RepeatModeler2 for automated genomic discovery of transposable element families. Proc. Natl. Acad. Sci. U. S. A. 117:9451–9457.

Frederick JR, Petri WA Jr. 2005. Roles for the galactose-/N-acetylgalactosamine-binding lectin of Entamoeba in parasite virulence and differentiation. Glycobiology 15:53R-59R.

Furukawa A, Nakada-Tsukui K, Nozaki T. 2012. Novel transmembrane receptor involved in phagosome transport of lysozymes and β-hexosaminidase in the enteric protozoan Entamoeba histolytica. PLoS Pathog. 8:e1002539.

Furukawa A, Nakada-Tsukui K, Nozaki T. 2013. Cysteine protease-binding protein family 6 mediates the trafficking of amylases to phagosomes in the Enteric protozoan Entamoeba histolytica. Infect. Immun. 81:1820–1829.

Grabherr MG, Haas BJ, Yassour M, Levin JZ, Thompson DA, Amit I, Adiconis X, Fan L, Raychowdhury R, Zeng Q, et al. 2011. Full-length transcriptome assembly from RNA-Seq data without a reference genome. Nat. Biotechnol. 29:644–652.

Hart AJ, Ginzburg S, Xu MS, Fisher CR, Rahmatpour N, Mitton JB, Paul R, Wegrzyn JL. 2020. EnTAP: Bringing faster and smarter functional annotation to non-model eukaryotic transcriptomes. Mol. Ecol. Resour. 20:591–604.

Hernandes-Alejandro M, Calixto-Gálvez M, López-Reyes I, Salas-Casas A, Cázares-Ápatiga J, Orozco E, Rodríguez MA. 2013. The small GTPase EhRabB of Entamoeba histolytica is differentially expressed during phagocytosis. Parasitol. Res. 112:1631– 1640.

Irmer H, Tillack M, Biller L, Handal G, Leippe M, Roeder T, Tannich E, Bruchhaus I. 2009. Major cysteine peptidases of Entamoeba histolytica are required for aggregation and digestion of erythrocytes but are dispensable for phagocytosis and cytopathogenicity. Mol. Microbiol. 72:658–667.

Jackman SD, Coombe L, Chu J, Warren RL, Vandervalk BP, Yeo S, Xue Z, Mohamadi H, Bohlmann J, Jones SJM, et al. 2018. Tigmint: correcting assembly errors using linked reads from large molecules. BMC Bioinformatics 19:393.

Jinatham V, Popluechai S, Clark CG, Gentekaki E. 2019. Entamoeba chiangraiensis n. sp. (Amoebozoa: Entamoebidae) isolated from the gut of Asian swamp eel (Monopterus albus) in northern Thailand. Parasitology 146:1719–1724.

Kato K, Tachibana H. 2022. Identification of multiple domains of Entamoeba histolytica intermediate subunit lectin-1 with hemolytic and cytotoxic activities. Int. J. Mol. Sci. 23:7700.

Katoh K, Standley DM. 2013. MAFFT multiple sequence alignment software version 7: improvements in performance and usability. Mol. Biol. Evol. 30:772–780.

Kawano T, Imada M, Chamavit P, Kobayashi S, Hashimoto T, Nozaki T. 2017. Genetic diversity of Entamoeba: Novel ribosomal lineages from cockroaches. PLoS One 12:e0185233.

Kawano-Sugaya T, Izumiyama S, Yanagawa Y, Saito-Nakano Y, Watanabe K, Kobayashi S, Nakada-Tsukui K, Nozaki T. 2020. Near-chromosome level genome assembly reveals ploidy diversity and plasticity in the intestinal protozoan parasite Entamoeba histolytica. BMC Genomics 21:813.

Keilwagen J, Hartung F, Grau J. 2019. GeMoMa: Homology-Based Gene Prediction Utilizing Intron Position Conservation and RNA-seq Data. Methods Mol. Biol. 1962:161–177.

Kim D, Paggi JM, Park C, Bennett C, Salzberg SL. 2019. Graph-based genome alignment and genotyping with HISAT2 and HISAT-genotype. Nat. Biotechnol. 37:907–915.

Kriventseva EV, Kuznetsov D, Tegenfeldt F, Manni M, Dias R, Simão FA, Zdobnov EM. 2019. OrthoDB v10: sampling the diversity of animal, plant, fungal, protist, bacterial and viral genomes for evolutionary and functional annotations of orthologs. Nucleic Acids Res. 47:D807–D811.

Lamichhaney S, Berglund J, Almén MS, Maqbool K, Grabherr M, Martinez-Barrio A, Promerová M, Rubin C-J, Wang C, Zamani N, et al. 2015. Evolution of Darwin’s finches and their beaks revealed by genome sequencing. Nature 518:371–375.

Li H. 2016. Minimap and miniasm: fast mapping and de novo assembly for noisy long sequences. Bioinformatics 32:2103–2110.

Li H. 2018. Minimap2: pairwise alignment for nucleotide sequences. Bioinformatics 34:3094–3100.

Loftus B, Anderson I, Davies R, Alsmark UCM, Samuelson J, Amedeo P, Roncaglia P, Berriman M, Hirt RP, Mann BJ, et al. 2005. The genome of the protist parasite Entamoeba histolytica. Nature 433:865–868.

Lorenzi HA, Puiu D, Miller JR, Brinkac LM, Amedeo P, Hall N, Caler EV. 2010. New assembly, reannotation and analysis of the Entamoeba histolytica genome reveal new genomic features and protein content information. PLoS Negl. Trop. Dis. 4:e716.

Lugo-Huitrón R, Blanco-Ayala T, Ugalde-Muñiz P, Carrillo-Mora P, Pedraza-Chaverrí J, Silva-Adaya D, Maldonado PD, Torres I, Pinzón E, Ortiz-Islas E, et al. 2011. On the antioxidant properties of kynurenic acid: free radical scavenging activity and inhibition of oxidative stress. Neurotoxicol. Teratol. 33:538–547.

Malik A, Dalal V, Ankri S, Tomar S. 2019. Structural insights into Entamoeba histolytica arginase and structure-based identification of novel non-amino acid based inhibitors as potential antiamoebic molecules. FEBS J. 286:4135–4155.

Marumo K, Nakada-Tsukui K, Tomii K, Nozaki T. 2014. Ligand heterogeneity of the cysteine protease binding protein family in the parasitic protist Entamoeba histolytica. Int. J. Parasitol. 44:625–635.

Meerovitch E. 1958. A NEW HOST OF ENTAMOEBA INVADENS RODHAIN, 1934. Can. J. Zool. 36:423–427.

Mikheenko A, Prjibelski A, Saveliev V, Antipov D, Gurevich A. 2018. Versatile genome assembly evaluation with QUAST-LG. Bioinformatics 34:i142–i150.

Min X, Feng M, Guan Y, Man S, Fu Y, Cheng X, Tachibana H. 2016. Evaluation of the C- Terminal Fragment of Entamoeba histolytica Gal/GalNAc Lectin Intermediate Subunit as a Vaccine Candidate against Amebic Liver Abscess. PLoS Negl. Trop. Dis. 10:e0004419.

Mitra BN, Saito-Nakano Y, Nakada-Tsukui K, Sato D, Nozaki T. 2007. Rab11B small GTPase regulates secretion of cysteine proteases in the enteric protozoan parasite Entamoeba histolytica. Cell. Microbiol. 9:2112–2125.

Nakada-Tsukui K, Marumo K, Nozaki T. 2020. A lysosomal hydrolase receptor, CPBF2, is associated with motility and invasion of the enteric protozoan parasite Entamoeba histolytica. Mol. Biochem. Parasitol. 239:111299.

Nakada-Tsukui K, Sekizuka T, Sato-Ebine E, Escueta-de Cadiz A, Ji D, Tomii K, Kuroda M, Nozaki T. 2018. AIG1 affects in vitro and in vivo virulence in clinical isolates of Entamoeba histolytica. PLoS Pathog. 14:e1006882.

Navarre C, Goffeau A. 2000. Membrane hyperpolarization and salt sensitivity induced by deletion of PMP3, a highly conserved small protein of yeast plasma membrane. EMBO J. 19:2515–2524.

Pertea G, Pertea M. 2020. GFF utilities: GffRead and GffCompare. F1000Res. 9:304.

Petri WA Jr, Chaudhry O, Haque R, Houpt E. 2006. Adherence-blocking vaccine for amebiasis. Arch. Med. Res. 37:288–291.

Price MN, Dehal PS, Arkin AP. 2010. FastTree 2--approximately maximum-likelihood trees for large alignments. PLoS One 5:e9490.

Quinlan AR, Hall IM. 2010. BEDTools: a flexible suite of utilities for comparing genomic features. Bioinformatics 26:841–842.

Ralston KS, Solga MD, Mackey-Lawrence NM, Somlata, Bhattacharya A, Petri WA. 2014. Trogocytosis by Entamoeba histolytica contributes to cell killing and tissue invasion. Nature 508:526–530.

Saito-Nakano Y, Loftus BJ, Hall N, Nozaki T. 2005. The diversity of Rab GTPases in Entamoeba histolytica. Exp. Parasitol. 110:244–252.

Saito-Nakano Y, Wahyuni R, Nakada-Tsukui K, Tomii K, Nozaki T. 2021. Rab7D small GTPase is involved in phago-, trogocytosis and cytoskeletal reorganization in the enteric protozoan Entamoeba histolytica. Cell. Microbiol. 23:e13267.

Saito-Nakano Y, Yasuda T, Nakada-Tsukui K, Leippe M, Nozaki T. 2004. Rab5-associated vacuoles play a unique role in phagocytosis of the enteric protozoan parasite Entamoeba histolytica. J. Biol. Chem. 279:49497–49507.

Sato D, Nakada-Tsukui K, Okada M, Nozaki T. 2006. Two cysteine protease inhibitors, EhICP1 and 2, localized in distinct compartments, negatively regulate secretion in Entamoeba histolytica. FEBS Lett. 580:5306–5312.

Scaglia M, Gatti S, Strosselli M, Grazioli V, Villa MR. 1983. Entamoeba moshkovskll (Tshalaia, 1941) Morpho-biological characterization of new strains isolated from the environment, and a review of the literature. Ann. Parasitol. Hum. Comp. 58:413–422.

Shiratori T, Ishida K-I. 2016. Entamoeba marina n. sp.; a New Species of Entamoeba Isolated from Tidal Flat Sediment of Iriomote Island, Okinawa, Japan. J. Eukaryot. Microbiol. 63:280–286.

Smith AJ, Barrett MT. 1915. The Parasite of Oral Endamebiasis, Endameba gingivalis (Gros). J. Parasitol. 1:159–174.

Suzek BE, Wang Y, Huang H, McGarvey PB, Wu CH, UniProt Consortium. 2015. UniRef clusters: a comprehensive and scalable alternative for improving sequence similarity searches. Bioinformatics 31:926–932.

Suzuki J, Kobayashi S, Murata R, Tajima H, Hashizaki F, Yanagawa Y, Takeuchi T. 2008. A survey of amoebic infections and differentiation of an Entamoeba histolytica-like variant (JSK2004) in nonhuman primates by a multiplex polymerase chain reaction. J. Zoo Wildl. Med. 39:370–379.

Tanaka M, Makiuchi T, Komiyama T, Shiina T, Osaki K, Tachibana H. 2019. Whole genome sequencing of Entamoeba nuttalli reveals mammalian host-related molecular signatures and a novel octapeptide-repeat surface protein. PLoS Negl. Trop. Dis. 13:e0007923.

Vaser R, Sović I, Nagarajan N, Šikić M. 2017. Fast and accurate de novo genome assembly from long uncorrected reads. Genome Res. 27:737–746.

Vera Alvarez R, Pongor LS, Mariño-Ramírez L, Landsman D. 2019. TPMCalculator: one- step software to quantify mRNA abundance of genomic features. Bioinformatics 35:1960–1962.

Verma K, Datta S. 2017. The Monomeric GTPase Rab35 Regulates Phagocytic Cup Formation and Phagosomal Maturation in Entamoeba histolytica. J. Biol. Chem. 292:4960–4975.

Walker BJ, Abeel T, Shea T, Priest M, Abouelliel A, Sakthikumar S, Cuomo CA, Zeng Q, Wortman J, Young SK, et al. 2014. Pilon: an integrated tool for comprehensive microbial variant detection and genome assembly improvement. PLoS One 9:e112963.

Welter BH, Powell RR, Leo M, Smith CM, Temesvari LA. 2005. A unique Rab GTPase, EhRabA, is involved in motility and polarization of Entamoeba histolytica cells. Mol. Biochem. Parasitol. 140:161–173.

Xu L, Dong Z, Fang L, Luo Y, Wei Z, Guo H, Zhang G, Gu YQ, Coleman-Derr D, Xia Q, et al. 2019. OrthoVenn2: a web server for whole-genome comparison and annotation of orthologous clusters across multiple species. Nucleic Acids Res. 47:W52–W58.

